# Antidiabetic drug therapy alleviates type 1 diabetes in mice by promoting pancreatic α-cell transdifferentiation

**DOI:** 10.1101/2020.08.03.234070

**Authors:** Dipak Sarnobat, Charlotte R Moffett, Neil Tanday, Frank Reimann, Fiona M Gribble, Peter R Flatt, Andrei I Tarasov

**Affiliations:** School of Biomedical Sciences, Ulster University, Cromore Road, Coleraine, BT52 1SA, UK; Metabolic Research Laboratories, Wellcome Trust-MRC Institute of Metabolic Science, Addenbrooke’s Hospital, Cambridge, CB2 0QQ, UK

**Keywords:** GLP-1 signalling, α/β-cell transdifferentiation, type 1 diabetes, apoptosis, glucagon

## Abstract

Gut incretins, glucagon-like peptide-1 (GLP-1) and glucose-dependent insulinotropic peptide (GIP), enhance secretion of insulin in a glucose-dependent manner, predominantly by elevating cytosolic levels of cAMP in pancreatic β-cells. Successful targeting of the incretin pathway by several drugs, however, suggests the antidiabetic mechanism is likely to span beyond the acute effect on hormone secretion and include, for instance, stimulation of β-cell growth and/or proliferation. Likewise, the antidiabetic action of kidney sodium-glucose linked transporter-2 (SGLT-2) inhibitors exceeds simple increase glucose excretion. Potential reasons for these ‘added benefits’ may lie in the long-term effects of these signals on developmental aspects of pancreatic islet cells. In this work, we explored if the incretin mimetics or SGLT-2 inhibitors can affect the size of the islet α- or β-cell compartments, under the condition of β-cell stress.

To that end, we utilised mice expressing YFP specifically in pancreatic α-cells, in which we modelled type 1 diabetes by injecting streptozotocin, followed by a 10-day administration of liraglutide, sitagliptin or dapagliflozin.

We observed an onset of diabetic phenotype, which was partially reversed by the administration of the antidiabetic drugs. The mechanism for the reversal included induction of β-cell proliferation, due to a decrease in β-cell apoptosis and, for the incretin mimetics, transdifferentiation of α-cells into β-cells.

Our data therefore emphasize the role of chronic incretin signalling in induction of α-/β-cell transdifferentiation. We conclude that incretin peptides may act directly on islet cells, making use of the endogenous local sites of ‘ectopic’ expression, whereas SGLT-2 inhibitors work via protecting β-cells from chronic hyperglycaemia.

**Graphical abstract:** 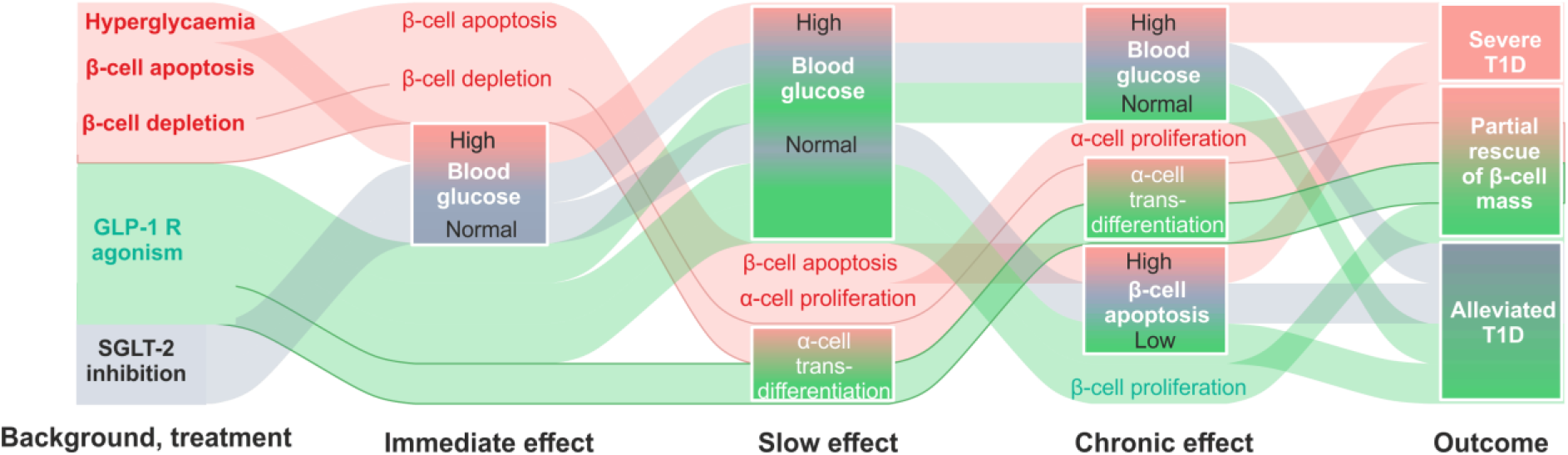

## Introduction

Independently of its aetiology, severe diabetes is associated with dysfunction of pancreatic islets of Langerhans, thereby compromising the hormonal regulation of blood glucose levels [1–3]. A home for cells of multiple types, islets release two key antagonising hormones, insulin (secreted by β-cells) and glucagon (α-cells), in concert, to ensure glucose clearance or recruitment into the systemic circulation, respectively. Notably, different islet cell populations derive from a common progenitor but undergo several stages of highly specific commitment, limiting their physiological role [4, 5]. An extreme β-cell-stress, however, has been reported to alter this pattern, by inducing a switch in the fate of pancreatic α- [6, 7], δ- [8] and non-islet pancreatic [9] cells. Relying on the islet cell plasticity, this phenomenon may represent a mechanism for potential regeneration of the β-cell mass upon its diabetes-induced loss [5].

The second largest islet cell subpopulation, α-cells have been conventionally viewed as the most appropriate cellular source for the β-cell regeneration [6, 10, 11]. This reputation has been further strengthened with recent studies that progressed from the therapeutically induced gain of β-cell properties by α-cells [12–15] to the ‘full-scale’ α-cell/β-cell transdifferentiation [7, 15–18]. The precise mechanism underlying α- to β-cell transition is, still unclear [11, 19] and may be functionally linked, for instance, to increased role of pancreatic GLP-1 signalling observed under substantial β-cell loss [1,20, 21].

Glucagon and GLP-1 derive from a common precursor, proglucagon, the products of two different prohormone convertases, PC2 and PC1/3, believed to be differentially expressed in pancreas or gut/brain proglucagon-expressing cells, respectively [1, 20]. Apart from being highly expressed in its ‘native’ L-cells, upon a β-cells loss, GLP-1 has also been reported in pancreatic α-cells, alongside glucagon [21]. This pattern arguably increases the local levels of the active forms of GLP-1 (7-36-amide and 7-37), whose systemic bioavailability is limited by the activity of a ubiquitous enzyme dipeptidyl peptidase-4 (DPP-4). The latter renders GLP-1 inactive (9-36) by cleaving the peptide bond between Ala^8^ and Glu^9^. Notably, the active form of GLP-1 that has been previously linked to the transdifferentiation between α- and β-cells [7, 22].

In this work, we examined the impact of a highly effective [1,23, 24] DPP-4 resistant synthetic GLP-1 analogue liraglutide and a DPP-4 inhibitor sitagliptin on α-cell proliferation and plasticity, in a mouse model. Alongside the two incretin mimetics we used a SGLT-2 inhibitor dapagliflozin [24, 25], to account for the glucose-lowering effect unrelated to the elevation of cytosolic cAMP or any other signalling mediated through GLP-1 receptor [26]. The present study was undertaken in mice bearing an inducible fluorescent label in α-cells (Glu^CreERT2^; ROSA26e-YFP) that were repeatedly treated with low doses of STZ to induce apoptosis in β-cells, which is expected to provide a critical signal to compensate for the β-cell loss.

## Materials and methods

### Animals

All experiments carried out under the UK Animals (Scientific Procedures) Act 1986 and EU Directive 2010/63EU were approved by the University of Ulster Animal Welfare and Ethical Review Body. Animals were maintained in an environmentally controlled laboratory at 22±2°C with a 12h dark and light cycle and given *ad libitum* access to standard rodent diet (10% fat, 30% protein and 60% carbohydrate; Trouw Nutrition, Northwich, UK) and water.

### Generation of Glu^CreERT2^;ROSA26-eYFP mice

Nine-week old Glu^CreERT2^;ROSA26-eYFP transgenic mice were used to perform all studies. An original colony, developed on the C57Bl/6 background at the University of Cambridge [27], was subsequently transferred to the animal facility at Ulster University and genotyped to assess Cre-ERT2 and ROSA26eYFP gene expression (Table 1). Three days prior to streptozotocin (STZ) dosing, mice were injected with tamoxifen (i.p. 7mg/mouse) to activate the tissue-specific expression of yellow fluorescent (YFP) in pancreatic islet α-cells (Figure 1A).

**Figure 1.**
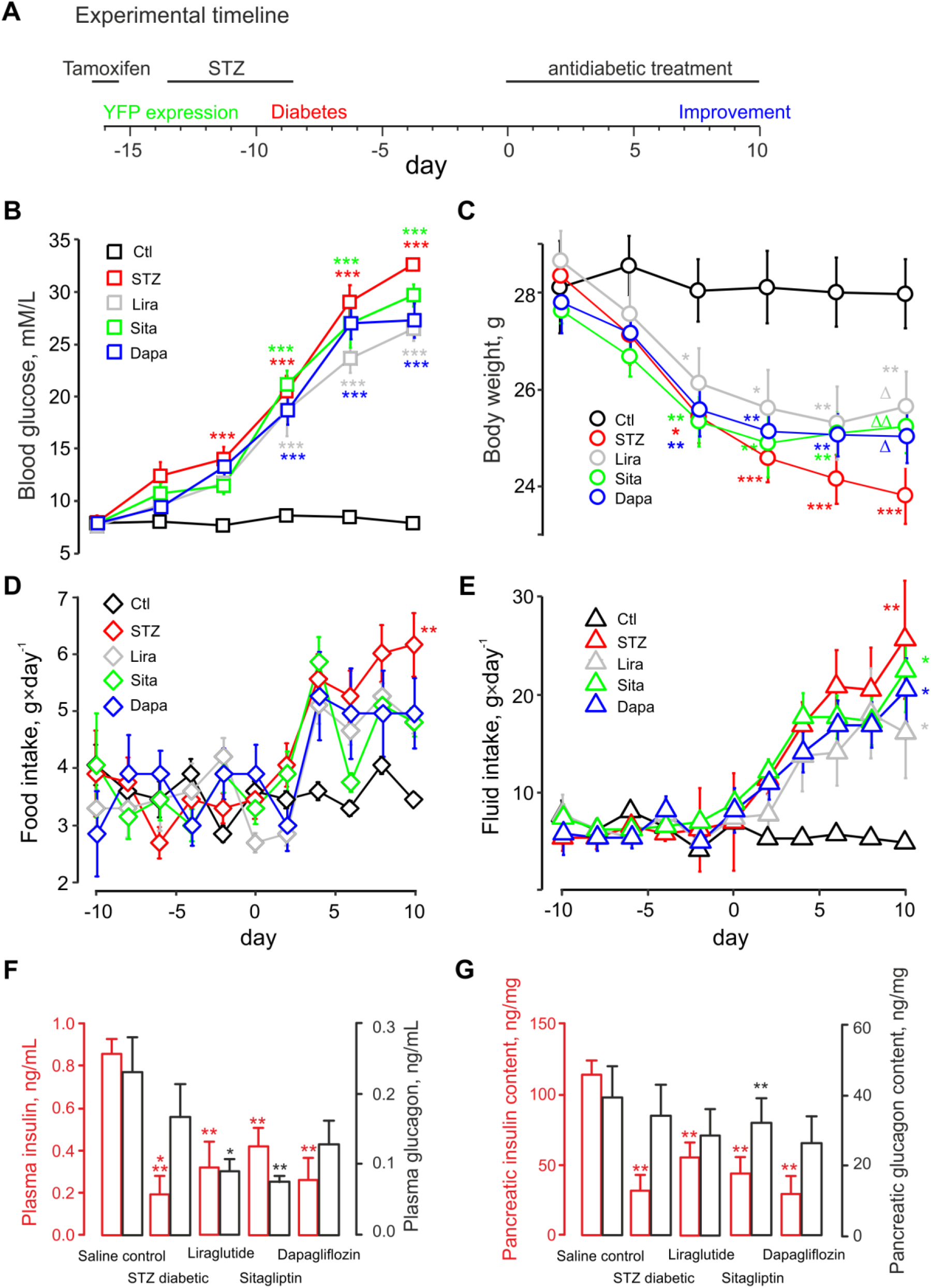
Liraglutide, sitagliptin and dapagliflozin partially rescue the diabetic phenotype of the streptozotocin-treated mice. ***A:*** Experimental timeline. Antidiabetic treatment starts on day 0. Tamoxifen if fed to the animals 16 days prior to that to induce the tissue-specific expression of YFP in α-cells. STZ is administered to model type 1 diabetes for four days, 9 days before the start of the treatment. The ability of the latter to improve the diabetic phenotype is is then assayed. ***B, C, D, E:*** Fasting blood glucose **(*B*)**, body weight **(*C*)**, food **(*D*)** and fluid ***(E)*** intake of Glu^CreERT2^;ROSA26-eYFP mice, following STZ treatment and the administration of antidiabetic drugs, as indicated, for groups of n=6 mice each. ‘STZ’, streptozotocin; ‘Sita’, sitagliptin; ‘Dapa’, dapagliflozin; ‘Lira’, liraglutide; ‘Ctl’, saline control. B-E ***F:*** plasma insulin (red) and glucagon (black). ***G:*** pancreatic insulin *(red)* and glucagon *(black)* content. ***F, G*** measurements were done on day 10, in different groups of mice, as indicated. *p<0.05, **p<0.01 vs saline control group. *p<0.05, **p<0.01 and ***p<0.001 compared to saline control group. Δp<0.05, ΔΔp<0.01 compared to the STZ group.

**Table 1.**
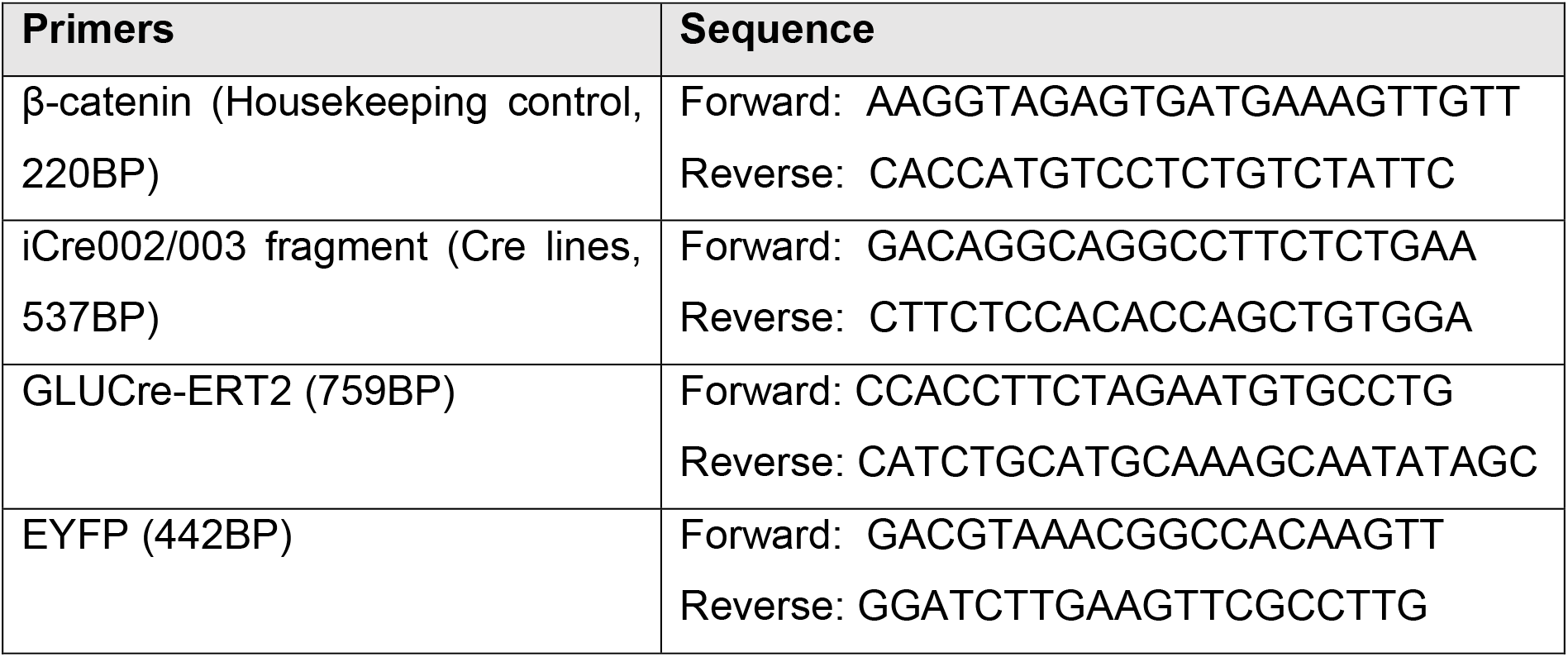
Primer sequence for PCR genotyping of Glu^CreERT2^;ROSA26-eYFP mice

### Diabetes model and antidiabetic medications

To develop insulin-deficient diabetes [21], 3 days after the induction of YFP expression, mice underwent a 5-day course of injections with low dose of STZ (50 mg/kg body weight daily, i.p.) (Figure 1A), dissolved in 0.1 M sodium citrate buffer (pH 4.5). The control group (n=6) received injections with the saline vehicle. The animals that underwent STZ injections were then divided into 4 groups (n=6) and treated with injections of saline vehicle, liraglutide (25 nmol/kg, intraperitoneal, 9:00 and 17:00 every day), sitagliptin (50 mg/kg, oral, once a day) or dapagliflozin (1 mg/kg, oral, once a daily) for 10 successive days. Cumulative food and fluid intake, body weight and blood glucose were assessed every 4 days. Non-fasting plasma insulin and glucagon were determined at the termination of the study (day 10).

### Blood glucose and hormone measurements

Blood samples were collected from the tail vein of animals into ice-chilled heparin-coated microcentrifuge tubes. Blood glucose was measured using a portable Ascencia meter (Bayer Healthcare, Newbury, Berkshire, UK). For plasma insulin and glucagon, blood was collected in chilled fluoride/heparin-coated tubes (Sarstedt, Numbrecht, Germany) and centrifuged using a Beckman microcentrifuge (Beckman Instruments, Galway, Ireland) for 10 minutes at 12,000 rpm. Plasma was extracted and stored at −20°C. For hormone determination from tissues, samples underwent acid-ethanol extraction (HCl: 1.5% v/v, ethanol: 75% v/v, H2O:23.5% v/v). Insulin concentrations were subsequently assessed by an in-house radioimmunoassay [28]. Plasma glucagon, pancreatic glucagon and GLP-1 content were measured using glucagon ELISA (EZGLU-30K, Merck Millipore), or RIA kit (250-tubes GL-32K, Millipore, USA) and GLP-1 ELISA kit Active (EGLP-35K, Millipore, MA, USA), respectively.

### Islet isolation and culturing

The mice were sacrificed by cervical dislocation. Islets were isolated using collagenase digestion [29]. A group of freshly isolated islets was directly embedded in agar (for immunohistochemistry) whereas the rest were cultured in RPMI-1640 medium containing 11.1 mM/L glucose and 0.3 g/L L-glutamine, supplemented with 10% fetal calf serum, 100 IU/mL penicillin, and 0.1 g/L streptomycin at 37°C for 72 hrs, in a fully humidified atmosphere with 5% CO2. To mimic diabetic conditions, the medium was supplemented with a mixture of cytokine factors (300 U/mL IL-1B, 300 U/mL IFNγ, 40 u/mL TNFα), as well as 0.25 mM sodium palmitate and 25 mM glucose [3, 21, 30]. Liraglutide was replaced every 12h, whereas other media components were changed every 24h.

### Preparation for immunostaining

After 72 h of culturing, islets were pelleted at the unit gravity, resuspended into 10% PFA and incubated for 10 min at 22°C. The fixed islets were then immobilised in warm 2% agarose gel. The samples were wrapped in a filter paper, placed into tissue cassette, processed (TP1020, Leica Microsystems, Nussloch, Germany) and embedded in paraffin wax [31]. 5 μm-thick sections were then immunostained for insulin and YFP.

### Immunohistochemistry and imaging

Following the removal of pancreatic tissue, samples were cut longitudinally and fixed with 4% PFA for 48 hr at 4°C. Fixed tissues were embedded and processed for antibody staining as described [21]. In brief, tissue sections (7μm) were blocked with 2% BSA, incubated with respective primary antibodies overnight at 4°C, and, subsequently, with appropriate conjugated secondary antibodies (Table 2). To stain nuclei, a final incubation was carried out at 37°C with 300 nM DAPI [Sigma-Aldrich, D9542]. To assess cell proliferation and/or apoptosis, co-staining of mouse anti-insulin (1:1000; Abcam, ab6995) or guinea pig anti-glucagon (PCA2/4, 1:200; raised in-house) with rabbit anti-Ki-67 (1:200; Abcam ab15580) or TUNEL reaction mixture (Roche Diagnostics Ltd, UK) was used. YFP tracing the α-cell lineage was detected by with a rabbit or goat anti-GFP (1:1000; Abcam, ab6556 or ab5450, respectively) (Table 2). The slides were imaged on an Olympus BX51 microscope, equipped with a 40x/1.3 objective. The multichannel fluorescence was visualised using DAPI (excitation 350 nm/emission 440 nm), FITC (488/515) and TRITC (594/610) filters and a DP70 camera controlled by Cell^F^ software (Olympus). Images were analysed using ImageJ software. All counts were determined in a blinded manner with 25-150 islets analysed per treatment group, as indicated in the figure legends.

**Table 2.** Primary and secondary antibodies used for immunohistochemistry

| Primary antibody | Dilution | Source |
| --- | --- | --- |
| Mouse anti-insulin | 1:1000 | Abcam, ab6995 |
| Guinea pig anti-glucagon | 1:200 | Raised in-house: PCA2/4 |
| Rabbit (goat) anti-GFP | 1:1000 | Abcam, ab6556 (ab5450) |
| Rabbit anti-Ki-67 | 1:200 | Abcam, ab15580 |
| TUNEL enzyme | 1:10 | Sigma Aldrich 11684795910 |
| Rabbit anti-GLP-1 (XJIC8) | 1:200 | in-house [20], specific for total GLP-1 |
| Secondary antibody | Dilution | Source |
| Goat anti-mouse | 1:400 | Alexa Fluor 488, Invitrogen, UK |
| Goat anti-mouse | 1:400 | Alexa Fluor 594, Invitrogen, UK |
| Goat anti-guinea pig | 1:400 | Alexa Fluor 488, Invitrogen, UK |
| Goat anti-guinea pig | 1:400 | Alexa Fluor 594, Invitrogen, UK |
| Goat anti-rabbit | 1:400 | Alexa Fluor 488, Invitrogen, UK |
| Donkey anti-rabbit | 1:500 | Alexa Fluor 594, Invitrogen, UK |
| Donkey anti-goat | 1:400 | Alexa Fluor 488, Invitrogen, UK |

### Data analysis and statistics

Statistical analysis was performed using GraphPad PRISM 5.0 or R [32]. Values are expressed as mean±SD, apart from the animal data (Figure 1) where the values are expressed as mean±SE. Comparative analyses between experimental groups were carried out using Student’s unpaired t-test or (for multiple groups) a one-way ANOVA with Bonferroni’s post-hoc test. The difference between groups was considered significant if P<0.05.

## Results

### STZ-induced type 1 diabetic phenotype is partially alleviated by the antidiabetic drugs

STZ treatment resulted in a progressive diabetic phenotype in the experimental animals, which was manifested in elevated levels of blood glucose (Figure 1B). Fasting glucose increased in the STZ-treated mice from 7.9±0.7 mM (end of the STZ treatment) to 32.2±0.8 mM 19 days afterwards (7.5±0.4 and 7.8±0.6 mM, respectively, in the control group). 10-day administration of liraglutide, sitagliptin or dapagliflozin had no significant impact on glycaemia (23.2±2.2, 29.9±0.8, 27.1 ±3.4 mM, respectively, p<0.05 vs control) (Figure 1A). The elevation in the resting glycaemia strongly correlated with the decrease in the body weight (Figure 1B). Body weight decreased from 28.2±0.6 g at the end of the STZ treatment to 23.8±0.5g 20 days afterwards, representing a loss off ca.15% (28.7±0.9 and 27.8±0.8 mM, respectively, in the control group, p<0.05) (Figure 1B). 10-day administration of the antidiabetic drugs mildly alleviated the weight loss (25.6±0.7, 25.3±0.5, 24.9±0.6 mM, for liraglutide, sitagliptin and dapagliflozin, respectively, p<0.05) (Figure 2B).

**Figure 2.**
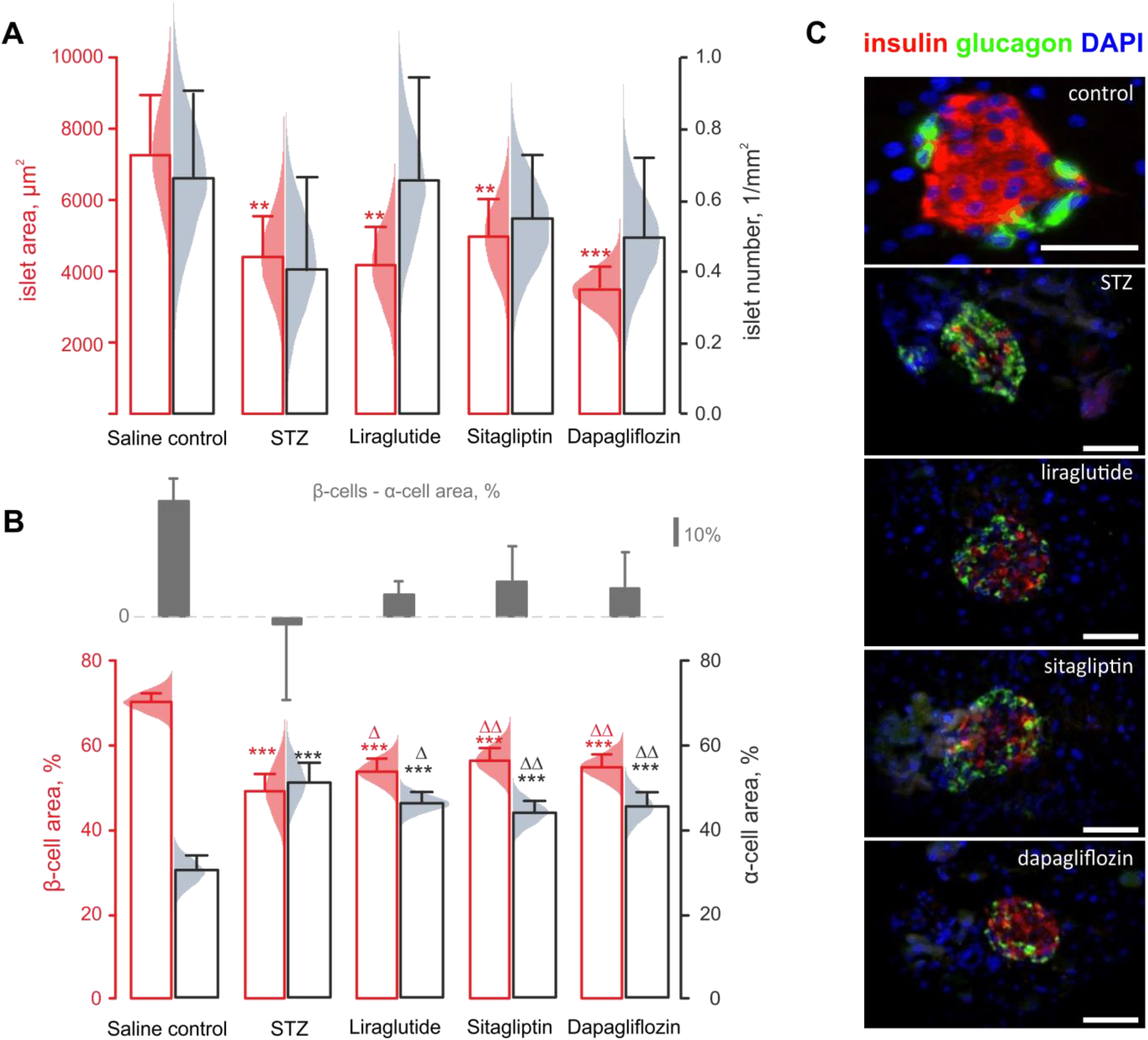
Diabetic phenotype is associated with changes in the islet composition. *A:* Islet area (red, n=150 islets from 6 mice) and number (black, n=150 islets from 6 mice). *B:* β-(red, n=150 islets from 6 mice) and α-cell (black, n=150 islets from 6 mice) percentage among the islet cells in response to the administration of STZ to Glu^CreERT2^;ROSA26-eYFP.mice and subsequent treatment with antidiabetic drugs, as indicated. Grey bars in *B* represent the net difference in the percentage of β- and α-cells. *C:* Representative immunostaining of mouse pancreatic sections for DAPI (blue), glucagon (green) and insulin (red). **p<0.01 and ***p<0.001 compared to saline control group. Δp<0.05 and ΔΔp<0.01 compared to streptozotocin treated group. Scale bars: 50μm.

The STZ treatment started affecting the intake of fluid or food by the mice only 11 days post cessation (day 2, Figure 1D,E), which coincided with the blood glucose increase over 15 mM in the four STZ-treated groups (Figure 1B). The intake of both fluid and food intake progressively elevated from that point, in the STZ-treated group (Figure 1D,E). Neither of the treatments was efficient in attenuating the fluid intake (Figure 1E), corresponding well to the lack of the reversal in hyperglycaemia (Figure 1B). However the highly variable data on food intake suggests that liraglutide, sitagliptin or dapagliflozin were able to interfere with that (intake on day 10: 4.8±0.2 4.8±0.2 5.0±0.6, respectively, ns vs 3.5±0.1 in the control group) (Figure 1D).

The STZ treatment resulted in a substantial decrease in non-fasting terminal plasma insulin levels, measured on day 10 (0.19±0.09 vs 0.87±0.05 ng/mL in STZ treated and control groups, respectively), with glucagon levels remaining practically unaffected (0.15±0.06 vs 0.21±0.07 ng/mL respectively, n.s.) (Figure 1F). None of the antidiabetic drugs was able to elevate insulin levels significantly (Figure 1F). At the same time, liraglutide and sitagliptin but not dapagliflozin treatment resulted in a significant decrease of plasma glucagon levels, on the STZ-treatment background (0.09±0.02 and 0.08±0.01 vs 0.23±0.05 ng/mL in the control group) (Figure 1F).

In line with the effect on plasma hormone levels (Figure 1F), STZ substantially decreased pancreatic content of insulin (33.28±10.94 vs 115.2±10.03 nM/mg of tissue in control, p<0.05), without any significant effect on the glucagon content (Figure 1G). The effect of the treatments on the insulin or glucagon content was not significant (Figure 1G).

### The alleviation of the diabetic phenotype is mediated via changes in the islet composition

The observations (Figure 1B-G) agreed well with the decrease in the average cross-section area of islets isolated from the Glu^CreERT2^;ROSA26-eYFP mice, treated with STZ, again with the antidiabetic drugs having no effect (red in Figure 2A,C). At the same time, we were unable to detect any significant alteration in the islet numbers per mm^2^, within any of the groups (black in Figure 2A), possibly due to substantial variability of this characteristic.

Compared to the control group, all STZ-treated groups showed a significant reduction in the relative β-cell area (red/insulin+ in Figure 2B,C) and, respectively, an increase in the relative α-cell area (black/glucagon+ in Figure 2B,C). Remarkably, liraglutide, sitagliptin and dapagliflozin partially rescued the effects of the STZ treatment on β- and α-cell fractions, resulting in small but significant differences in the percentage of β-cells (53±1%, 56±1%, 54±1%, respectively, vs 48±3% in STZ mice) and α-cells (46±1%, 44±1%, 45±1% vs 50±2% in STZ mice) (Figure 2B,C).

### Antidiabetic drugs decrease apoptosis and increase the proliferation of β-cells

Despite being able to induce β-cell necrosis when administered at a single high dose, STZ has been shown to induce β-cell apoptosis, when used in small repeated doses [33]. In line with this report, we observed a nine-fold (3.5±0.3 vs 0.4±0.1% in control mice) increase in the percentage of β-cells expressing an apoptosis marker, TUNEL, in mice treated with STZ (red in Figure 3A). The β-cell apoptosis was substantially (dapagliflozin) or practically completely (liraglutide, sitagliptin) reversed by the antidiabetic drugs (red, Figure 3A). Finally, none of the STZ-treated groups displayed any changes in α-cell apoptosis frequency (black, Figure 3A).

**Figure 3.**
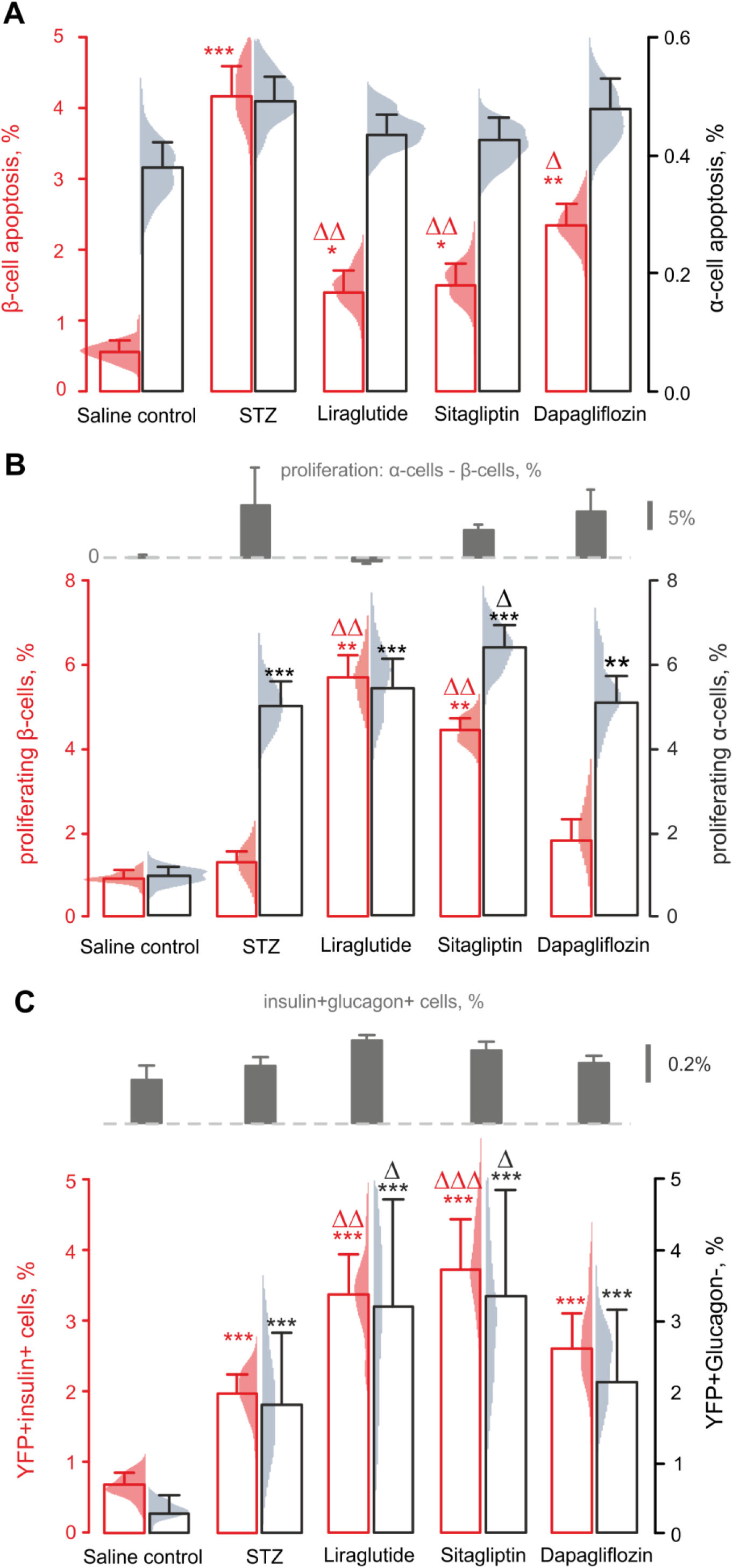
Antidiabetic drugs decrease β-cell apoptosis, increase proliferation and transdifferentiation from α-cells. ***A, B:*** Percentage of β-cells (red, n=60 islets from 6 mice) and α-cells (black, n=60 islets from 6 mice) undergoing apoptosis **(*A*)**, as determined by TUNEL staining, or proliferation ***(B)***, Ki-67 staining, in response to the administration of STZ to Glu^CreERT2^;ROSA26-eYFP.mice and subsequent treatment with antidiabetic drugs, as indicated. Grey bars in ***B*** represent the net difference in the proliferating fractions of α- and β-cells. ***C*:** Trans-differentiation of YFP+ cells within Glu^CreERT2^;ROSA26-eYFP.mice. The YFP expression, originally specifically induced in α-cells, was detectable within non-α-cells (***C***, black, n=60 islets from 6 mice) and β-cells (***C***, red, n=60 islets from 60 mice) after to the administration of STZ and subsequent anti-diabetic treatment. In addition, double-positive (insulin+glucagon+) cells were detectable (***C***, grey bars, n=60 islets from 6 mice). *p<0.05, **p<0.01 and ***p<0.001 compared to saline control group. Δp<0.05 and ΔΔp<0.01, ΔΔΔp<0.001 compared to STZ-treated group.

The pro-apoptotic effect of the STZ treatment was not associated with any increase in the percentage of proliferating β-cells, probed via Ki-67 staining (red in Figure 3B). It did however produce a 5-fold increase in the fraction of proliferating α-cells (black in Figure 3B), with neither of the antidiabetic treatments attenuating the α-cell proliferation (black in Figure 3B). At the same time, liraglutide and sitagliptin significantly increased the proliferation frequency of β-cells, post the STZ treatment (red in Figure 3B).

### Long-term administration of antidiabetic drugs induces α-/β-cell transdifferentiation

In the Glu^CreERT2^; ROSA26-eYFP mice, expression of YFP can be induced specifically in pancreatic α-cells. When co-detected with anti-glucagon antibodies, 26 days post induction of the targeted YFP expression, the islets from the control mouse group had a small percentage of YFP+ cells that did not express glucagon (0.3±0.1%), which was increased 6-fold after the STZ treatment (1.8±0.4%) and further potentiated by liraglutide and sitagliptin (3.2±0.6%, 3.4±0.6%, respectively) but not dapagliflozin (2.2±1%) (black in Figure 3C).

Whilst fairly low in the control animals (0.8±0.1%), the percentage of the YFP+insulin+ cells in the STZ-treated mice was 2.5 times higher (1.9±0.2%) than in the control group (red in Figure 3C, red). Just as in case of YFP+glucagon-cells, liraglutide and sitagliptin (3.3±0.2% and 3.8±0.3%, respectively) further potentiated the commitment of the YFP+ cells towards the insulin lineage, whereas dapagliflozin had no statistically significant (2.6±0.2%) effect (red, Figure 3C). The differentiation of YFP+ α-cells towards the insulin lineage was reflected in the increased percentage of bi-hormonal cells, upon the incretin mimetic or DPP-4 inhibitor treatment (gray bars above Figure 3C). The relative size of this small cell subpopulation was practically unaffected by the STZ treatment alone (0.35±0.02% vs 0.26±0.05% in the control group), however, liraglutide (0.49±0.03%) and sitagliptin (0.43±0.02%) and, to a lesser extent, dapagliflozin (0.37±0.01%) administration substantially expanded the population of bi-hormonal cells (gray bars, Figure 3C).

### Insulin mimetics induce α-/β-cell transdifferentiation ex vivo

Liraglutide, sitagliptin and dapagliflozin have multiple reported targets within the body. To test whether the expansion of the insulin+YFP+ cell population in the mouse model could result from a direct effect of these medications on pancreatic islets, we isolated the islets of Langerhans from Glu^CreERT2^; ROSA26-eYFP mice and modelled the diabetic/apoptotic conditions in tissue culture for 72 hours, with(out) rescue by liraglutide, sitagliptin and dapagliflozin (Figure 4A). The *ex vivo* culturing resulted in an increase in basal fraction of YFP+insulin+ cells (2.0± 0.05%), perhaps reflecting the technological differences between the *in vivo* and *ex vivo* models. The basal percentage of YFP+insulin+ cells was unaffected by conditions mimicking diabetes/apoptosis (high glucose, high palmitate, IL-1B, IFNγ, TNFα) (Figure 4B) thereby contrasting the *in vivo* data (Figure 3C). However, just like in the case of *in vivo* administration, the *ex vivo* exposure to liraglutide or sitagliptin was able to substantially expand the fraction of YFP+insulin+ cells (7.7±0.2% and 5.8±0.2%, respectively, vs 2.0±0.5% in the control group, p<0.05), whereas the effect of dapagliflozin was not significant (2.5±0.3%).

**Figure 4.**
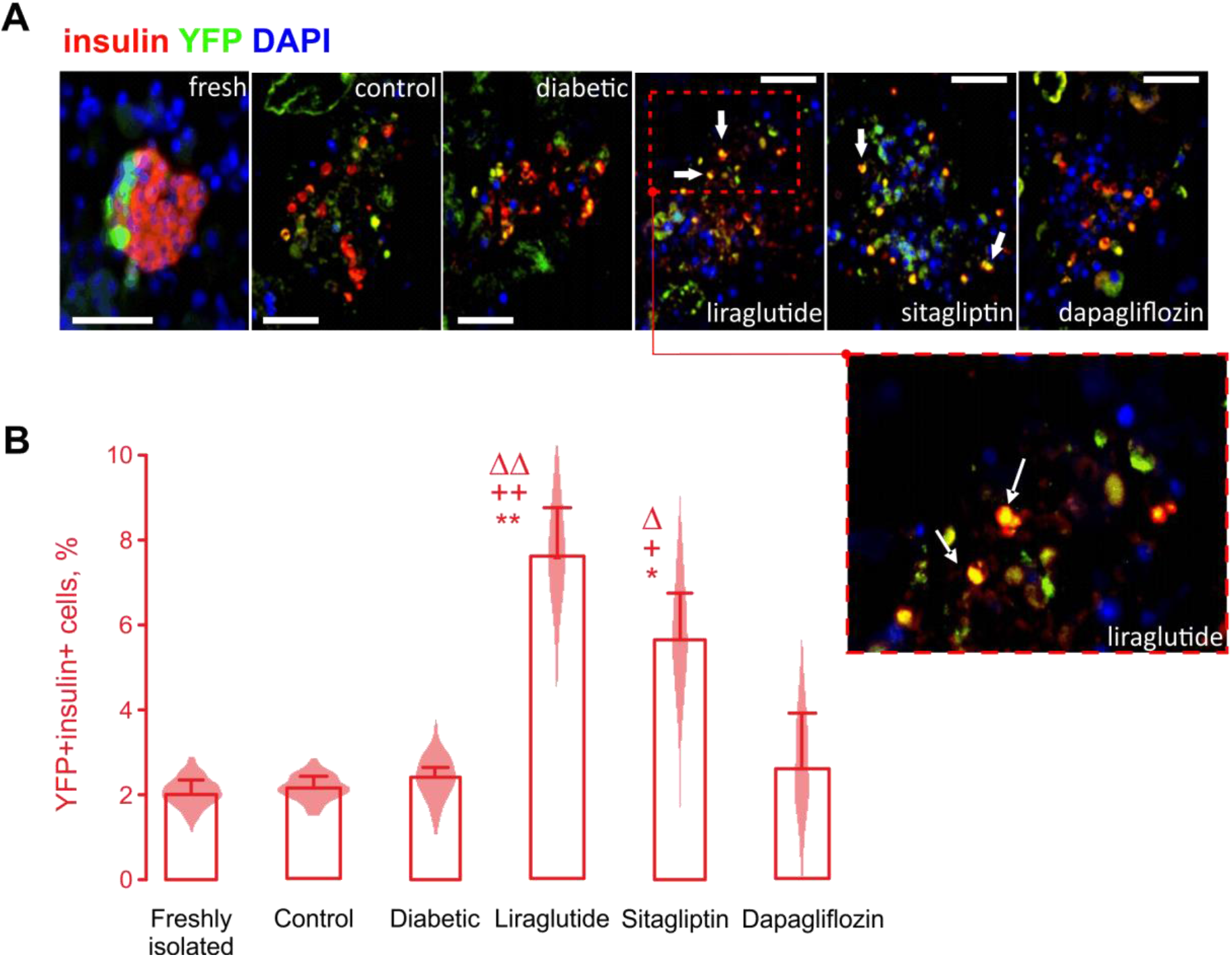
Incretin mimetics but not the SGLT-2 inhibitor enhance α-/β-transdifferentiation of isolated islet cells. *A:* Representative images showing immunostaining for DAPI (blue), YFP (green) and insulin (red) in islets freshly isolated from Glu^CreERT2^;ROSA26-eYFP mice as well as islets cultured for 72 h in RPMI medium with different supplements (as indicated, detailed in Islet isolation and culturing). Scale bars: 50μm. *B:* Percentage of YFP+insulin+ cells reflecting the recruitment of α-cells along the β-cell lineage, in response to the culturing conditions (as in *A*) for n=25 islets from 3 mice per group. *p<0.05 and **p<0.01 compared to freshly isolated group; +p<0.05 and ++p<0.01 compared to the control RPMI-1640 media; Δp<0.05 and ΔΔp<0.01 compared to cytokine mixture (diabetic control).

### Streptozotocin stimulates expression of GLP-1 by α-cells

Whilst liraglutide targets GLP-1 receptors expressed on pancreatic islet cells, sitagliptin elevates the circulating levels of incretins by inhibiting DPP-4, an enzyme that renders the incretins inactive. Sitagliptin therefore is unlikely to have any insulinotropic or antidiabetic effect in the absence of the systemic incretins, a condition used in the *ex vivo* experiments (Figure 4). We dissected the potential mechanism of the expansion of YFP+insulin+ cell population, upon chronic exposure to sitagliptin, by quantifying the levels of GLP-1 within islets isolated from the Glu^CreERT2^;ROSA26-eYFP mice (Figure 5A,B). GLP-1 was practically undetectable within YFP+ and YFP-cell populations, in the control group of animals (Figure 5A; apparent staining in the β-cell pool is the background fluorescence). The administration of STZ induced a significant expansion of the YFP+ cells (Figure 5A, red in Figure 5B), in line with the increase in α-cell percentage and proliferation rate (Figure 2B, Figure 3B). Furthermore, STZ elevated the numbers of GLP-1+ cells, almost exclusively among the YFP+ cells (Figure 5A). In line with that, the GLP-1 content in the islets was dramatically increased, after the STZ treatment (black in Figure 5B).

**Figure 5.**
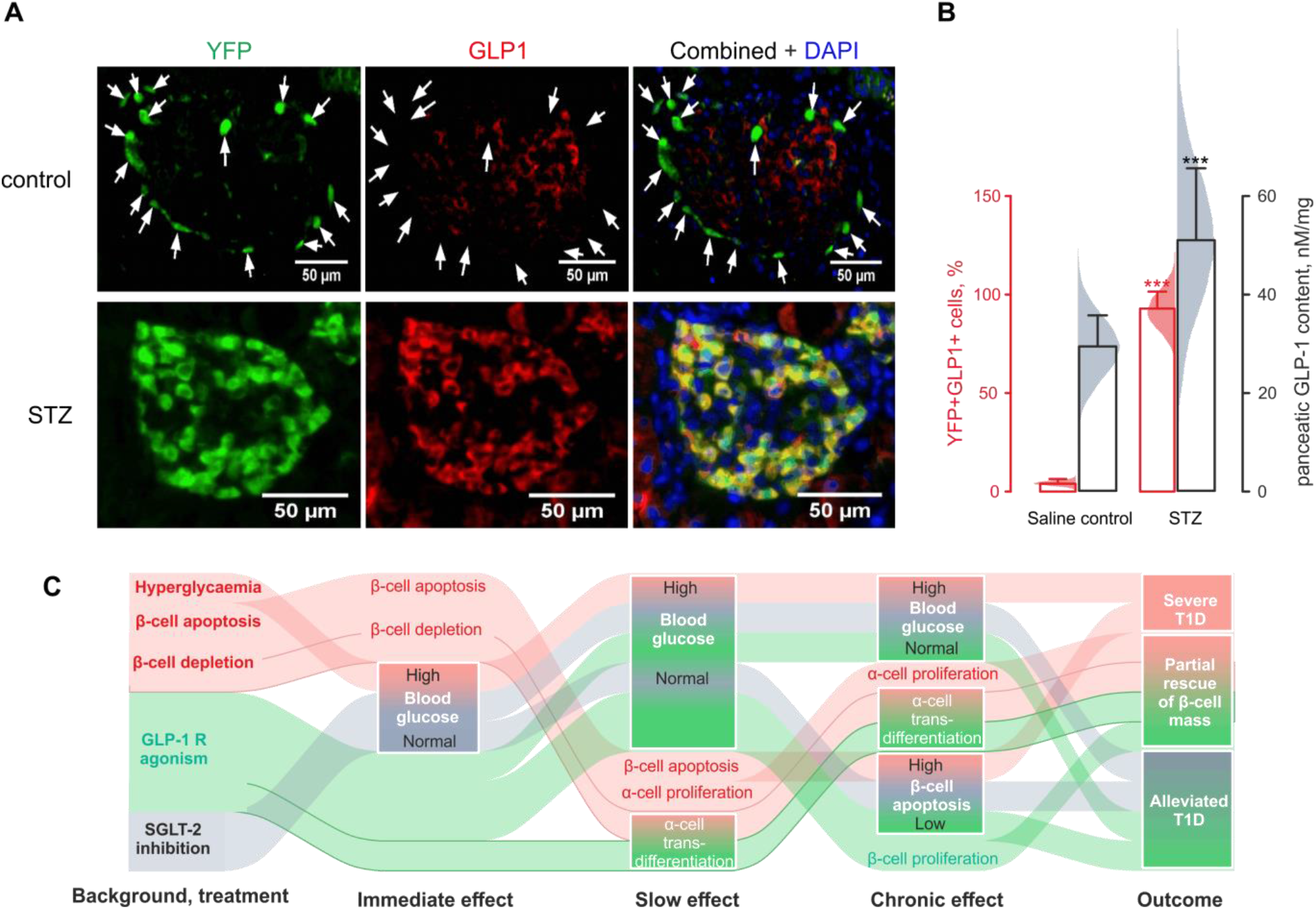
Potential mechanism: STZ treatment increases expression of GLP-1 by α-cells. *A:* Representative images showing immunostaining of pancreases from control and STZ-treated mice for DAPI (blue), YFP (green) and GLP-1 (red). Arrows indicate GLP-1^-^/YFP+ α-cells from saline control group. Scale bars: 50μm. *B:* Effect of STZ on immuno-expression of GLP-1 by YFP-labelled α-cells (red, n=150 islets from 6 mice) and pancreatic GLP-1 content measured using ELISA (black, n=150 islets from 6 mice). **p<0.01 and ***p<0.001 compared to saline control group. *C:* schematic of the proposed mechanism.

## Discussion

We report the plasticity within the cell populations of pancreatic islets of Langerhans. In our hands, pancreatic β-cell subpopulation was partially replenished via reprogramming of α-cells, following an induced apoptotic β-cell injury.

### Mouse phenotype, islet mass and morphology

In line with previous observations [34], this study demonstrated the development of a reliable diabetic phenotype, upon repeated injections of small doses of STZ: a week after the end of the 5-day injection course, the animals displayed stably elevated glycaemia (Figure 1B). This correlated with a reduced body weight (Figure 1C), indicating the impaired ability of metabolising glucose by the body or increased dehydration due to polyuria. The phenotype was worsening throughout all the remaining 12 days of the study, with the three antidiabetic drugs being able to only partially rescue it, as blood glucose, body weight and blood insulin remained substantially altered. At the same time, treatment with liraglutide and sitagliptin reduced blood glucagon levels, reflecting an acute attenuation mechanism working directly on α-cells [35] or indirectly, via an elevation of somatostatin secretion from neighbouring δ-cells [36]. The decrease in circulating glucagon is unlikely to reflect the recruitment of the α-cell population into trans-differentiation process (Figure 1D), as the pool of ‘resting’ α-cells is quite large, in rodent islets [37, 38]. Multiple low doses of STZ, in our hands, affected the islet size and cellular composition, as has been reported earlier [21]. Furthermore, liraglutide, sitagliptin and dapagliflozin increased the relative β-cell and reduced α-cell area, consistent with previous reports [1, 21, 30]. We examined whether the increase in the β-cell mass is derived from improved survival, enhanced proliferation or rather a *de novo* production. The STZ treatment had no effect on β-cell proliferation but, as expected, dramatically enhanced β-cell apoptosis (Figure 3A) [21]. All three antidiabetic drugs attenuated the apoptosis of β-cells (Figure 3A), however they differed in their impact on β-cell proliferation. Both liraglutide and sitagliptin substantially increased the proliferation of β-cells, in line with [21,23, 25], whereas dapagliflozin was ineffective in this respect (Figure 3B). Of note, neither liraglutide, nor sitagliptin attenuated the proliferation of α-cells, which was induced by the STZ treatment (Figure 3B).

### Expression of insulin by YFP-tagged α-cells

The STZ-induced severe β-cell injury *per se* resulted in a detectable expression of insulin and a loss of expression of glucagon by cells bearing the YFP label (Figure 3C), which was originally specifically targeted to α-cell population. Liraglutide and sitagliptin but not dapagliflozin further increased the co-expression of insulin and YFP on a per-cell basis, suggesting a role for the incretin signalling in regulating α-/β-cell transdifferentiation (Figure 3C).

A fraction of α-cells has been reported to start producing insulin, without an immediate loss of glucagon expression, thereby storing two hormones at the same time in the same cell [6, 7]. These bi-hormonal cells would eventually become deficient in glucagon and progress into fully mature β-cells [6]. In the present study, a small but detectable number of bi-hormonal cells was evident after the STZ treatment, which was further increased by liraglutide and sitagliptin (Figure 3C).

### Potentiation of cell differentiation and proliferation by incretins

The role of non-Glut family glucose transporters in the glucose sensing by α-cells is a matter of intensive research [26]. Consistent with our findings, inhibition of SGLT-2 was reported to enhance glucagon secretion from isolated islets [26], whilst inhibition of SGLT-1 attenuated the secretion of the hormone [26]. Long-term administration of dapagliflozin has been reported to expand the β-cell mass by inhibiting apoptosis [39], which agrees well with our findings (Figure 3A).

Although GLP-1 expression by pancreatic α-cells in response to the depletion of the β-cell numbers has been reported previously [7, 20, 21], the mechanism linking the two phenomena remains unclear. Speculatively, it is likely to include a yet unidentified β-cell injury signal targeting the neighbouring α-cells, resulting in an activation of proconvertase PC1/3 and hence GLP-1 production [40]. As such, local GLP-1 may be (in)directly involved in the regeneration of β-cells [7]. Potential mechanisms of GLP-1-mediated α-cell transdifferentiation include PI3K/AKT/FOXO1 pathway [22], fibroblast growth factor 21 [7] or epidermal growth factor [41] signalling. However, it remains to be determined to what extent the detectable GLP-1 reflects active GLP-1(7-37) and GLP-1(7-36-amide), or the corresponding 1-37/1-36-amide forms, which lack GLP1R-activity.

### Consensus model

The three antidiabetic drugs administered to the experimental Glu^CreERT2^;ROSA26-eYFP mice interacted with STZ-induced apoptosis of the islet β-cells as well as its direct consequences, reduction of the β-cell mass and hyperglycaemia (Figure 5C). Long-acting GLP-1 receptor agonist liraglutide and DPP-4 inhibitor sitagliptin have a delayed absorption kinetics (8 hours) and a long half-life (12-13 hours) [42]. SGLT-2 inhibitor dapagliflozin has a similar half-life in humans [43], which is however 2.5 times shorter in rodents [44]; dapagliflozin reaches the maximal plasma concentration much faster (within 1.5 hours [45]). The two GLP-1 mimetics are therefore likely to counteract hyperglycaemia on the ‘slow’ timescale of ca.10 hours whereas dapagliflozin has a ‘fast’ mode of action (1...10 h) (Figure 5C). Although none of the 10-day long treatments was able to reverse the STZ-induced T1D hyperglycaemic phenotype, all three medications significantly attenuated β-cell apoptosis (Figure 3A, Figure 5C), possibly by reducing the peak plasma glucose concentrations [46]. On the ‘chronic’ timescale (several days), the depletion of β-cells by the STZ pre-treatment (Figure 3A) enhanced the proliferation of α-cells (Figure 3B) as well as α-/β-cell transdifferentiation (Figure 3C). The GLP-1 receptor agonism upregulated both processes in α-cells (Figure 3C), which resulted in a partial recovery of the β-cell mass (Figure 5C, accented). Although the latter could have been further facilitated by GLP-1-dependent potentiation of β-cell proliferation (Figure 3B), the proliferation may also be hypothetically attributed to the newly trans-differentiated α-cells.

We therefore believe that the mechanism of the upregulation of α-cell transdifferentiation includes a direct effect of the incretin mimetics on pancreatic islet cells, as was proven via experiments on isolated islets (Figure 4). Whereas the GLP-1 receptor agonist targets the islet cells directly, the DPP-4 inhibitor is likely to rely on the endogenous production of the incretin by islet α-cells, via a hypothetical switch from PC2 to PC1/3. A highly contested finding for non-diabetic native mouse islets, GLP-1 expression in α-cells is likely to be induced by the diabetic conditions (Figure 5), in line with recent human data [47]. At the same time, the *ex vivo* (non-diabetic) effect of sitagliptin on the α-cell transdifferentiation (Figure 4B) is at odds with this mechanism and is therefore likely to be linked to an induction of islet GLP-1 expression in response to the chronic (72 h) culture conditions.

### Conclusions

Despite their short *in vivo* half-life and the fast nature of signalling via the membrane receptors, incretin peptides are also involved in mediating chronic (days) processes within the body. Those include energy metabolism, that underlies such features of the incretin drugs as cardioprotection, as well as cell differentiation and proliferation. The availability of synthetic long-acting GLP-1 receptor agonists as albiglutide and dulaglutide prompts for studies of the mechanisms underlying this long-timescale signals, with a special focus on cell proliferation and plasticity in human islets where, possibly due to the age-associated factors, both processes are believed to be attenuated.

## Acknowledgements

These studies were supported in part by young investigator award to RCM and Ulster University Vice-Chancellor Research Studentship award to DS. Research in the Reimann/Gribble laboratory is currently funded by the Wellcome Trust (106262/Z/14/Z and 106263/Z/14/Z) and the MRC (MRC_MC_UU_12012/3).

## Abbreviations

T1D: type 1 diabetes
GLP-1: glucagon-like peptide-1
SGLT: sodium-glucose linked transporter 2
YFP: yellow fluorescent protein
STZ: streptozotocin
DPP-4: dipeptidyl peptidase 4
GIP: glucose-dependent insulinotropic peptide
TNF: tumour necrosis factor
IFN: interferon
IL: interleukin
FITC: fluorescein isothiocyanate
DAPI: 4’,6-diamidino-2-phenylindole
TUNEL: terminal deoxynucleotidyl transferase dUTP nick end labelling

## References

[1] R.C. Moffett, S. Patterson, N. Irwin, P.R. Flatt, Positive effects of GLP-1 receptor activation with liraglutide on pancreatic islet morphology and metabolic control in C57BL/KsJ db/db mice with degenerative diabetes, Diabetes/metabolism research and reviews 31(3) (2015) 248–255.

[2] S. Vasu, N.H. McClenaghan, P.R. Flatt, Molecular mechanisms of toxicity and cell damage by chemicals in a human pancreatic beta cell line, 1.1 B4, Pancreas 45(9) (2016) 1320–1329.

[3] S. Vasu, R.C. Moffett, N.H. McClenaghan, P.R. Flatt, Differential molecular and cellular responses of GLP-1 secreting L-cells and pancreatic alpha cells to glucotoxicity and lipotoxicity, Experimental cell research 336(1) (2015) 100–108.

[4] T. van der Meulen, M.O. Huising, The role of transcription factors in the transdifferentiation of pancreatic islet cells, Journal of molecular endocrinology 54(2) (2015) R103.

[5] S. Puri, A.E. Folias, M. Hebrok, Plasticity and dedifferentiation within the pancreas: development, homeostasis, and disease, Cell stem cell 16(1) (2015) 18–31.

[6] F. Thorel, V. Népote, I. Avril, K. Kohno, R. Desgraz, S. Chera, P.L. Herrera, Conversion of adult pancreatic α-cells to β-cells after extreme β-cell loss, Nature 464(7292) (2010) 1149–1154.

[7] S.-H. Lee, E. Hao, D. Scharp, F. Levine, Insulin acts as a repressive factor to inhibit the ability of PAR2 to induce islet cell transdifferentiation, Islets 10(6) (2018) 201–212.

[8] S. Chera, D. Baronnier, L. Ghila, V. Cigliola, J.N. Jensen, G. Gu, K. Furuyama, F. Thorel, F.M. Gribble, F. Reimann, Diabetes recovery by age-dependent conversion of pancreatic δ-cells into insulin producers, Nature 514(7523) (2014) 503–507.

[9] M. Solar, C. Cardalda, I. Houbracken, M. Martín, M.A. Maestro, N. De Medts, X. Xu, V. Grau, H. Heimberg, L. Bouwens, Pancreatic exocrine duct cells give rise to insulin-producing β cells during embryogenesis but not after birth, Developmental cell 17(6) (2009) 849–860.

[10] Y. Cheng, H. Kang, J. Shen, H. Hao, J. Liu, Y. Guo, Y. Mu, W. Han, Beta-cell regeneration from vimentin+/MafB+ cells after STZ-induced extreme beta-cell ablation, Scientific reports 5 (2015) 11703.

[11] V. Stanojevic, J.F. Habener, Evolving function and potential of pancreatic alpha cells, Best Practice & Research Clinical Endocrinology & Metabolism 29(6) (2015) 859–871.

[12] P. Collombat, X. Xu, P. Ravassard, B. Sosa-Pineda, S. Dussaud, N. Billestrup, O.D. Madsen, P. Serup, H. Heimberg, A. Mansouri, The ectopic expression of Pax4 in the mouse pancreas converts progenitor cells into α and subsequently β cells, Cell 138(3) (2009) 449–462.

[13] Y.-P. Yang, F. Thorel, D.F. Boyer, P.L. Herrera, C.V. Wright, Context-specific α-to-β-cell reprogramming by forced Pdx1 expression, Genes & development 25(16) (2011) 1680–1685.

[14] C.L. Wilcox, N.A. Terry, E.R. Walp, R.A. Lee, C.L. May, Pancreatic α-cell specific deletion of mouse Arx leads to α-cell identity loss, PloS one 8(6) (2013).

[15] J. Lu, P.L. Herrera, C. Carreira, R. Bonnavion, C. Seigne, A. Calender, P. Bertolino, C.X. Zhang, α Cell–Specific Men1 Ablation Triggers the Transdifferentiation of Glucagon-Expressing Cells and Insulinoma Development, Gastroenterology 138(5) (2010) 1954–1965. e8.

[16] A.M. Ackermann, N.G. Moss, K.H. Kaestner, GABA and artesunate do not induce pancreatic α-to-β cell transdifferentiation in vivo, Cell metabolism 28(5) (2018) 787–792. e3.

[17] N. Ben-Othman, A. Vieira, M. Courtney, F. Record, E. Gjernes, F. Avolio, B. Hadzic, N. Druelle, T. Napolitano, S. Navarro-Sanz, Long-term GABA administration induces alpha cell-mediated beta-like cell neogenesis, Cell 168(1-2) (2017) 73–85. e11.

[18] J. Li, T. Casteels, T. Frogne, C. Ingvorsen, C. Honore, M. Courtney, K.V. Huber, N. Schmitner, R.A. Kimmel, R.A. Romanov, Artemisinins target GABAA receptor signaling and impair α cell identity, Cell 168(1-2) (2017) 86–100. e15.

[19] J. Gromada, I. Franklin, C.B. Wollheim, α-Cells of the endocrine pancreas: 35 years of research but the enigma remains, Endocrine reviews 28(1) (2007) 84–116.

[20] R.C. Moffett, S. Vasu, B. Thorens, D.J. Drucker, P.R. Flatt, Incretin receptor null mice reveal key role of GLP-1 but not GIP in pancreatic beta cell adaptation to pregnancy, PloS one 9(6) (2014).

[21] S. Vasu, R.C. Moffett, B. Thorens, P.R. Flatt, Role of endogenous GLP-1 and GIP in beta cell compensatory responses to insulin resistance and cellular stress, PloS one 9(6) (2014).

[22] Z. Zhang, Y. Hu, N. Xu, W. Zhou, L. Yang, R. Chen, R. Yang, J. Sun, H. Chen, A New Way for Beta Cell Neogenesis: Transdifferentiation from Alpha Cells Induced by Glucagon-Like Peptide 1, Journal of diabetes research 2019 (2019).

[23] B.D. Green, P.R. Flatt, C.J. Bailey, Dipeptidyl peptidase IV (DPP IV) inhibitors: a newly emerging drug class for the treatment of type 2 diabetes, Diabetes and vascular disease research 3(3) (2006) 159–165.

[24] P. Millar, N. Pathak, V. Parthsarathy, A.J. Bjourson, M. O’Kane, V. Pathak, R.C. Moffett, P.R. Flatt, V.A. Gault, Metabolic and neuroprotective effects of dapagliflozin and liraglutide in diabetic mice, Journal of Endocrinology 234(3) (2017) 255–267.

[25] S.-i. Asahara, W. Ogawa, SGLT2 inhibitors and protection against pancreatic beta cell failure, Springer, 2019.

[26] T. Suga, O. Kikuchi, M. Kobayashi, S. Matsui, H. Yokota-Hashimoto, E. Wada, D. Kohno, T. Sasaki, K. Takeuchi, S. Kakizaki, SGLT1 in pancreatic α cells regulates glucagon secretion in mice, possibly explaining the distinct effects of SGLT2 inhibitors on plasma glucagon levels, Molecular metabolism 19 (2019) 1–12.

[27] J.R. Campbell, A. Martchenko, M.E. Sweeney, F. Michael, A. Psichas, F.M. Gribble, F. Reimann, P.L. Brubaker, Essential role of Munc18-1 in the regulation of glucagon-like peptide-1 secretion, https://tspace.library.utoronto.ca/handle/1807/98201.

[28] P. Flatt, C. Bailey, Abnormal plasma glucose and insulin responses in heterozygous lean (ob/+) mice, Diabetologia 20(5) (1981) 573–577.

[29] Y.H. Abdel-Wahab, F.P. O’Harte, H. Ratcliff, N.H. McClenaghan, C.R. Barnett, P.R. Flatt, Glycation of insulin in the islets of Langerhans of normal and diabetic animals, Diabetes 45(11) (1996) 1489–1496.

[30] S. Vasu, R. Moffett, J.T. McCluskey, M. Hamid, N. Irwin, P. Flatt, Beneficial effects of parenteral GLP-1 delivery by cell therapy in insulin-deficient streptozotocin diabetic mice, Gene therapy 20(11) (2013) 1077–1084.

[31] D. Bosco, M. Armanet, P. Morel, N. Niclauss, A. Sgroi, Y.D. Muller, L. Giovannoni, G. Parnaud, T. Berney, Unique arrangement of α-and β-cells in human islets of Langerhans, Diabetes 59(5) (2010) 1202–1210.

[32] R Development Core Team., R: A Language and Environment for Statistical Computing. <https://www.R-project.org/>, 2019).

[33] B.A. O’BRIEN, B.V. Harmon, D.P. Cameron, D.J. Allan, Beta-cell apoptosis is responsible for the development of IDDM in the multiple low-dose streptozotocin model, The Journal of pathology 178(2) (1996) 176–181.

[34] D. Sarnobat, R.C. Moffett, V.A. Gault, N. Tanday, F. Reimann, F.M. Gribble, P.R. Flatt, N. Irwin, Effects of long-acting GIP, xenin and oxyntomodulin peptide analogues on alpha-cell transdifferentiation in insulin-deficient diabetic GluCreERT2; ROSA26-eYFP mice, Peptides (2019) 170205.

[35] Y.Z. De Marinis, A. Salehi, C.E. Ward, Q. Zhang, F. Abdulkader, M. Bengtsson, O. Braha, M. Braun, R. Ramracheya, S. Amisten, A.M. Habib, Y. Moritoh, E. Zhang, F. Reimann, A.H. Rosengren, T. Shibasaki, F. Gribble, E. Renstrom, S. Seino, L. Eliasson, P. Rorsman, GLP-1 inhibits and adrenaline stimulates glucagon release by differential modulation of N-and L-type Ca2+ channel-dependent exocytosis, Cell metabolism 11(6) (2010) 543–53.

[36] A. Ørgaard, J.J. Holst, The role of somatostatin in GLP-1-induced inhibition of glucagon secretion in mice, Diabetologia 60(9) (2017) 1731–1739.

[37] A.I. Tarasov, J. Galvanovskis, O. Rorsman, A. Hamilton, E. Vergari, P.R. Johnson, F. Reimann, F.M. Ashcroft, P. Rorsman, Monitoring real-time hormone release kinetics via high-content 3-D imaging of compensatory endocytosis, Lab on a Chip 18(18) (2018) 2838–2848.

[38] A. Hamilton, Q. Zhang, A. Salehi, M. Willems, J.G. Knudsen, A.K. Ringgaard, C.E. Chapman, A. Gonzalez-Alvarez, N.C. Surdo, M. Zaccolo, D. Basco, P.R.V. Johnson, R. Ramracheya, G.A. Rutter, A. Galione, P. Rorsman, A.I. Tarasov, Adrenaline Stimulates Glucagon Secretion by Tpc2-Dependent Ca(2+) Mobilization From Acidic Stores in Pancreatic alpha-Cells, Diabetes 67(6) (2018) 1128–1139.

[39] A. Kanno, S.i. Asahara, M. Kawamura, A. Furubayashi, S. Tsuchiya, E. Suzuki, T. Takai, M. Koyanagi-Kimura, T. Matsuda, Y. Okada, Early administration of dapagliflozin preserves pancreatic β-cell mass through a legacy effect in a mouse model of type 2 diabetes, J Diabetes Invest 10(3) (2019) 577–590.

[40] N. Whalley, L. Pritchard, D. Smith, A. White, Processing of proglucagon to GLP-1 in pancreatic a-cells: is this a paracrine mechanism enabling GLP-1 to act on b-cells, J Endocrinol 211 (2011) 99–106.

[41] J. Fusco, X. Xiao, K. Prasadan, Q. Sheng, C. Chen, Y.-C. Ming, G. Gittes, GLP-1/Exendin-4 induces β-cell proliferation via the epidermal growth factor receptor, Scientific reports 7(1) (2017) 1–6.

[42] G.A. Herman, C. Stevens, K. Van Dyck, A. Bergman, B. Yi, M. De Smet, K. Snyder, D. Hilliard, M. Tanen, W. Tanaka, Pharmacokinetics and pharmacodynamics of sitagliptin, an inhibitor of dipeptidyl peptidase IV, in healthy subjects: results from two randomized, double-blind, placebo-controlled studies with single oral doses, Clinical Pharmacology & Therapeutics 78(6) (2005) 675–688.

[43] FARXIGA-dapagliflozin tablet, film coated, https://dailymed.nlm.nih.gov/dailymed/drugInfo.cfm?setid=72ad22ae-efe6-4cd6-a302-98aaee423d69 [accessed 15/05/2020].

[44] M. Obermeier, M. Yao, A. Khanna, B. Koplowitz, M. Zhu, W. Li, B. Komoroski, S. Kasichayanula, L. Discenza, W. Washburn, In vitro characterization and pharmacokinetics of dapagliflozin (BMS-512148), a potent sodium-glucose cotransporter type II inhibitor, in animals and humans, Drug metabolism and disposition 38(3) (2010) 405–414.

[45] G.S. Tirucherai, F. Lacreta, F.A. Ismat, W. Tang, D.W. Boulton, Pharmacokinetics and pharmacodynamics of dapagliflozin in children and adolescents with type 2 diabetes mellitus, Diabetes, Obesity and Metabolism 18(7) (2016) 678–684.

[46] J. Adam, R. Ramracheya, M.V. Chibalina, N. Ternette, A. Hamilton, A.I. Tarasov, Q. Zhang, E. Rebelato, N.J.G. Rorsman, R. Martin-del-Rio, A. Lewis, G. Ozkan, H.W. Do, P. Spegel, K. Saitoh, K. Kato, K. Igarashi, B.M. Kessler, C.W. Pugh, J. Tamarit-Rodriguez, H. Mulder, A. Clark, N. Frizzell, T. Soga, F.M. Ashcroft, A. Silver, P.J. Pollard, P. Rorsman, Fumarate Hydratase Deletion in Pancreatic beta Cells Leads to Progressive Diabetes, Cell Reports 20(13) (2017) 3135–3148.

[47] S.A. Campbell, D. Golec, M. Hubert, J. Johnson, N. Salamon, A. Barr, P.E. MacDonald, K. Philippaert, P.E. Light, Human islets contain a subpopulation of glucagon-like peptide-1 secreting α cells that is increased in type 2 diabetes, Molecular Metabolism (2020) 101014.

